# Membrane voltage fluctuations in human breast cancer cells

**DOI:** 10.1101/2021.12.20.473148

**Authors:** Peter Quicke, Yilin Sun, Mar Arias-Garcia, Corey D. Acker, Mustafa B. A. Djamgoz, Chris Bakal, Amanda J. Foust

## Abstract

Cancer cells feature a resting membrane potential (*V*_*m*_) that is depolarized compared to normal cells, and express active ionic conductances, which factor directly in their pathophysiological behavior. Despite similarities to ‘excitable’ tissues, relatively little is known about cancer cell *V*_*m*_ dynamics. With high-throughput, cellular-resolution *V*_*m*_ imaging, we characterized *V*_*m*_ fluctuations of hundreds of human triple-negative breast cancer MDA-MB-231 cells and compared to non-cancerous breast epithelial MCF-10A cells. By quantifying their Dynamic Electrical Signatures (DESs) through an unsupervised machine-learning protocol, we identified four classes ranging from “noisy” to “blinking/waving”. The *V*_*m*_ of MDA-MB-231 cells exhibited spontaneous, transient hyperpolarizations that were inhibited by the voltage-gated sodium channel blocker tetrodotoxin. The *V*_*m*_ of MCF-10A cells was comparatively static, but fluctuations increased following treatment with transforming growth factor-β1, a canonical inducer of the epithelial-to-mesenchymal transition. These data suggest that the ability to generate *V*_*m*_ fluctuations is acquired during transformation and may participate in oncogenesis.

## Introduction

All cells in the body exhibit a voltage difference (*V*_*m*_) across the plasma membrane and use this to regulate a wide range of functions such as gene expression, secretion, and whole-cell motility. Cellular *V*_*m*_ at rest varies both between and within cell types. Interestingly, whilst this is ca. −70 mV in mature ‘quiescent’ cells, including nerves and muscles, it is noticeably depolarized (*V*_*m*_ ~ −50 to −10 mV) in proliferating cells, including cancer cells and stem cells^1^,^2^.

*V*_*m*_ fluctuates dramatically, both spontaneously and in response to stimuli, in classically excitable tissues such as heart, muscle and nerve, which support the generation and conduction of action potentials. The resting *V*_*m*_ of several cell types has been shown to fluctuate^3^. These include cells with rhythmic activity e.g. neurons controlling respiration^4^, arterial vasomotion^5^, biological ‘clocks’^6^ and sleep^7^,^8^. Oscillations of *V*_*m*_ also manifest in pathophysiological situations such as epilepsy and neuronal degeneration, and can extend to network effects^9^,^10^.

In several carcinomas, functional expression of voltage-gated sodium channels (VGSCs) promotes the metastatic process^11^. Treating carcinoma cells in vitro with VGSC blockers partially suppresses 3D invasion^12^,^13^. The most specific inhibitor of VGSCs is tetrodotoxin (TTX), which blocks the channel by binding to a site within the channel pore when the channel is in the open state^14^. TTX reduces invasion in carcinoma cells in-vitro, and this effect is abolished by siRNA silencing of the VGSC Nav1.5 in-vivo^12,13,15,16^. Gradek et al. recently demonstrated that reduced expression of SIK1 induces Nav1.5 expression, invasion and the expression of EMT-associated transcription factor SNAl1^17^. However, as noted above, the steady-state resting *V*_*m*_ of human breast cancer cells relative to normal epithelia is strongly depolarized^18^. In the case of the MDA-MB-231 cells, derived from a highly aggressive triple-negative breast cancer, *V*_*m*_ rests between −40 mV to −20 mV^15^,^19,20^. The *V*_*m*_-dependent inactivation of VGSCs means that the majority of channels should be permanently inactivated at such depolarized membrane potentials and therefore insensitive to TTX. Nevertheless, TTX has been shown repeatedly to inhibit the invasiveness of these cells and several other carcinomas^11,15,19,21–24,^ potentially by blocking the persistent window current^19^,^25^.

Although the depolarization of resting *V*_*m*_ and the enriched VGSC expression in aggressive cancer cell lines are established (reviewed by^2^ and^26^), unlike classical excitable tissues (e.g., heart, muscle, nerve), comparatively little is known about cancer’s *V*_*m*_ dynamics. Studies utilizing multi-electrode arrays detected *V*_*m*_ fluctuations but could not to attribute them to individual cells^27^,^28^. Here in contrast we captured cellular-resolution, spatially resolved *V*_*m*_ dynamics in human breast cancer cells with a fast, electrochromic voltage-sensitive dye, enabling optical monitoring of *V*_*m*_ changes in hundreds of cells simultaneously. Through an unsupervised machine learning protocol, we classified and characterized the Dynamic Electrical Signatures (DESs) of the cellular *V*_*m*_ time series obtained with high-throughput imaging. A subset of MDA-MB-231 breast cancer cells exhibited hyperpolarizing “blinks” and “waves”, in contrast with the quiescent, static *V*_*m*_ of non-tumorigenic MCF-10A cells. Application of TTX suppressed the *V*_*m*_ fluctuations in MDA-MB-231 cells whilst treatment of MCF-10A cells with transforming growth factor-β1 (TGF-β), which stimulates the epithelial-to-mesenchymal transition (EMT), induced *V*_*m*_ fluctuations in these cells. Taken together, these data suggest that the ability to generate *V*_*m*_ fluctuations is acquired during the EMT and may participate in oncogenesis.

## Results

### Di-4-AN(F)EP(F)PTEA fluorescence ratio linearly reports change in *V*_*m*_

We imaged the membrane potential of cultured cell monolayers with extracellularly-applied di-4-AN(F)EP(F)PTEA^29^, a dye that inserts into the outer membrane leaflet, shifting its absorption and emission spectra as a function of membrane potential with sub-microsecond temporal fidelity. We sequentially excited the dye with blue and green light-emitting diodes (LEDs, Figure 1A,B; Supplementary Figure S1), taking the ratio of fluorescence excited by each color at each point in time and dividing by the baseline ratio (Δ*R/R*_0_). A change in *V*_*m*_ causes the fluorescence excited by each color to change in opposite directions (Figure 1C), amplifying the corresponding change in the ratio. The ratiometric imaging scheme also partially mitigates the confounds of uneven dye labeling, photobleaching decay, and mechanical motion^30^. This approach enabled us to image the dynamics of hundreds of human breast cancer cells simultaneously with cellular resolution.

**Figure 1.**
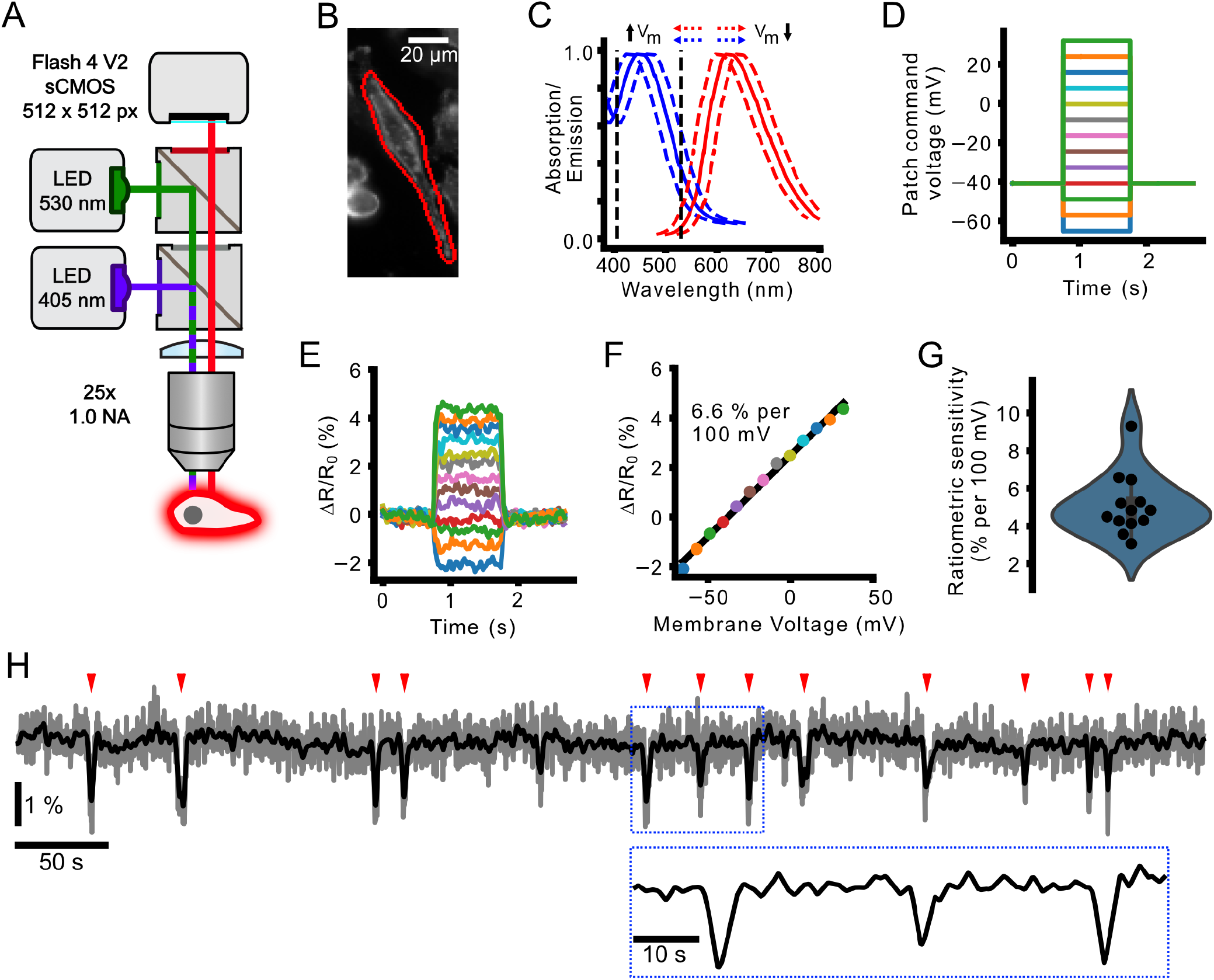
Electrochromic imaging of MDA-MB-231 cells with voltage dye di-4-AN(F)EP(F)PTEA. (A) Schematic of the widefield epifluorescence imaging system with two-color excitation. (B) MDA-MB-231 cell stained with di-4-AN(F)EP(F)PTEA with simultaneous voltage clamp (C) The blue and red lines show di-4-AN(F)EP(F)PTEA’s excitation and emission spectra^29^, respectively. Fluorescence is excited by blue and green LEDs on opposite sides of the excitation maximum (black vertical dashed lines). Decreasing or increasing *V*_*m*_ causes the excitation spectra to shift right or left, respectively. Decreasing *V*_*m*_ therefore causes reduced emission in response to blue channel excitation, and increased fluorescence with green channel excitation and vice versa for increasing *V*_*m*_. (D) Waveforms commanding the MDA-MB-231 whole-cell voltage clamp for calibration of the ratio of blue- to green-excited fluorescence with respect to the baseline (Δ*R/R*_0_). (E) Recorded Δ*R/R*_0_ signal in response to the injected *V*_*m*_ waveforms. (F) Average Δ*R/R*_0_ change with membrane voltage change and a linear fit to the data indicating the imaging sensitivity (% change in ratio per 100 mV membrane voltage change). (G) Measured sensitivities for different patched cells with a gaussian kernel estimate (blue envelope). (H) Example time course of a spontaneously active MDA-MB-231 cell displaying typically observed transient *V*_*m*_ hyperpolarisations (indicated by red ticks). Grey, unfiltered time course, black, gaussian filtered time course, sigma = 3 sampling points (0.6 s).

We first verified that the fluorescence ratio (Δ*R/R*_0_) linearly reported changes in *V*_*m*_. By imaging Di-4-AN(F)EP(F)PTEA fluorescence while stepping *V*_*m*_ through a range of values in whole-cell voltage clamp of MDA-MB-231 cells, we observed that the ratio of blue-to-green Di-4-AN(F)EP(F)PTEA fluorescence varied linearly with changes in *V*_*m*_ over a physiological range of *V*_*m*_ ranging from –60 to +30 mV. The slope of Δ*R/R*_0_ vs *V*_*m*_ showed an average sensitivity of 5.1 ± 0.43 % per 100 mV (mean ± standard error of the mean, (s.e.m.); n = 12 cells; Figure 1D-G). Furthermore, as expected, global depolarization of all cells on the coverslip by washing in a high-potassium (100 mM) extracellular solution also increased Δ*R/R*_0_ (Supplementary Figure S2). These results show that the ratio Δ*R/R*_0_ faithfully reports *V*_*m*_ changes in subsequent recordings of spontaneous *V*_*m*_ fluctuations (Figure 1H).

### Cellular-resolution membrane voltage fluctuations in MDA-MB-231 cells

A subset of MDA-MB-231 cells exhibited fluctuating *V*_*m*_. We imaged spontaneous *V*_*m*_ fluctuations at 5 frames/second in cultured monolayers of the highly aggressive triple-negative breast cancer cell line MDA-MB-231 (Figure 2A). 6.84% ± 0.97% (n=22 coverslips) of cells on each coverslip displayed a fluctuating *V*_*m*_ (Figure 2B,C). The great majority (>90%, 91/100) of high signal-to-noise ratio, large amplitude (>|1.5%| Δ*R/R*_0_) events were negative-going (“-VE”, Figure 2B). Therefore, subsequent analyses focus on the −VEs (Figure 2A, B). Most cells exhibited few or no fluctuations, but a subset of cells were highly active (Figure 2C, see also supplementary Movie M1). Among the active cells, events were detected at an average frequency of 2 ± 0.2 events/cell/1000 seconds (n= 20 coverslips, Figure 2D).

**Figure 2.**
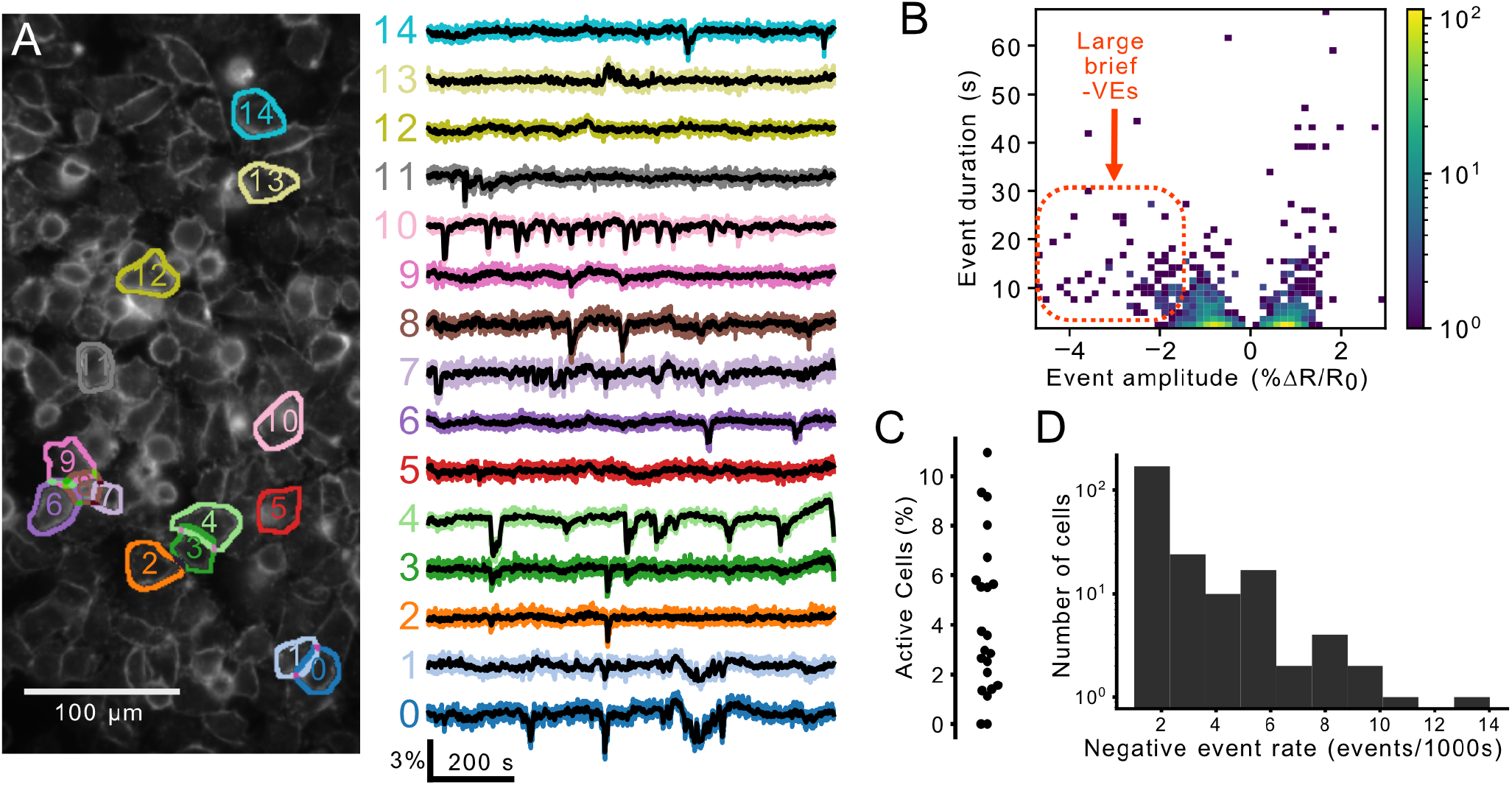
A subset of MDA-MB-231 cells exhibit *V*_*m*_ fluctuations (A) The ratio of Di-4-AN(F)EP(F)PTEA fluorescence (Δ*R/R*_0_) was imaged at 5 frames/second in cultured MDA-MB-231 cells. The Δ*R/R*_0_ time series extracted from the segmented cells (right) reveals heterogeneous *V*_*m*_ fluctuations consisting primarily of transient hyperpolarizations. (B) A log-scaled 2D histogram of event amplitude and duration. These fluctuations vary in their polarity (positive-going, “+VE” vs. negative-going “-VE”), amplitude, and duration. Large amplitude fluctuations were typically hyperpolarising (-VE amplitude) and short in duration. (C) Proportion of active cells for 20 technical replicates displaying the high variability in the proportion of active cells. (D) A log-scaled histogram of the mean −VE rate for all active cells (at least one −VE observed) showing the long-tailed distribution with a few highly active cells.

We then sought to describe the “Dynamic Electrical Signature” (DES) of individual cells based on their *V*_*m*_ imaging time series. A DES is a multi-parametric feature vector which captures various aspects of *V*_*m*_ fluctuations over time such as power spectral properties, successive differences, and entropy. A DES captures variations beyond the event-based metrics and can be used to classify patterns based on their similarity. To describe the DESs, we implemented an unsupervised machine learning pipeline (Figure 3). The Cellular DES Pipeline uses the Catch-22 algorithm^31^ to extract features from individual *V*_*m*_ traces. Feature extraction and hierarchical clustering on the “active” cellular time series, those in which *V*_*m*_ fluctuations were detected (287 out of 2993, Figure 3A-C), yielded silhouette coefficients indicating that the time series clustered into 3 or 4 classes (Figure 3D). Manual inspection of the time series revealed higher inner-cluster similarly with 4 clusters. We named the DES classes identified by the 4 cluster classification: small blinking (blinking-S), waving, noisy, and large blinking (blinking-L). Figure 4 displays exemplar time series from each class. Cells of the four classes were observed simultaneously in the imaged fields-of-view (FOVs). Together these four DES classes describe the temporal heterogeneity of *V*_*m*_ fluctuations in MDA-MB-231 cells.

**Figure 3.**
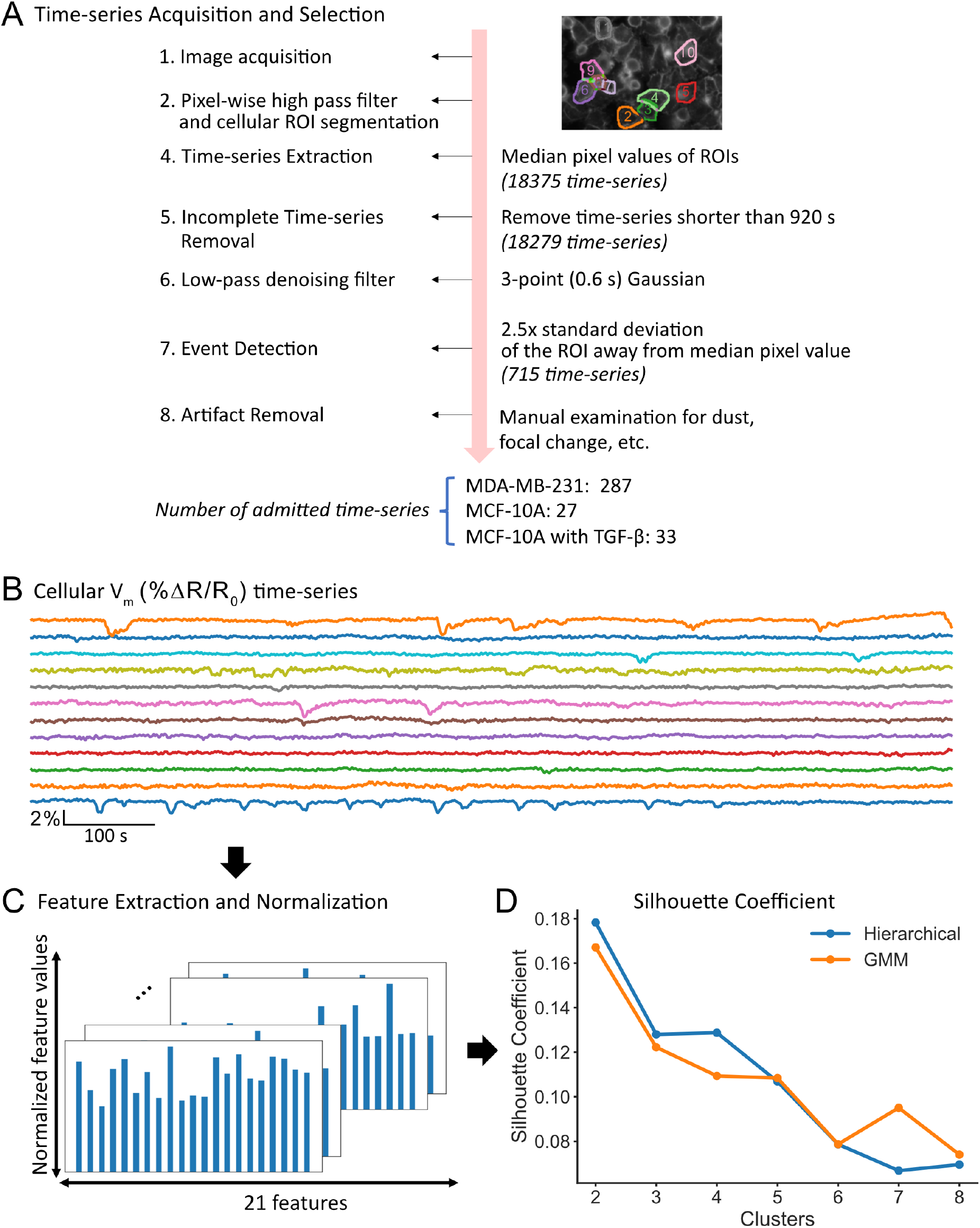
The Dynamic Electrical Signature (DES) of MDA-MB-231 cells clustered into four classes (A) Time series were admitted to the unsupervised machine learning clustering pipeline if their temporally filtered Δ*R/R*_0_ exhibited fluctuations > 2.5x of their within-ROI pixel standard deviation from the median pixel value, and if also free of imaging artefacts including floating dust, mechanical vibration and illumination edge effects. (B) Visual inspection of each cell’s time series finalized admission to the Cellular DES Pipeline. (C) 22 features relevant to time series temporal patterns were extracted from each time series using the Catch-22 algorithm^31^ and normalized with a Box Cox transformation relative to their values. (D) The silhouette coefficients for different cluster numbers were generated through hierarchical clustering (blue) or Gaussian Mixture Modeling (GMM, orange) on the normalized features.

**Figure 4.**
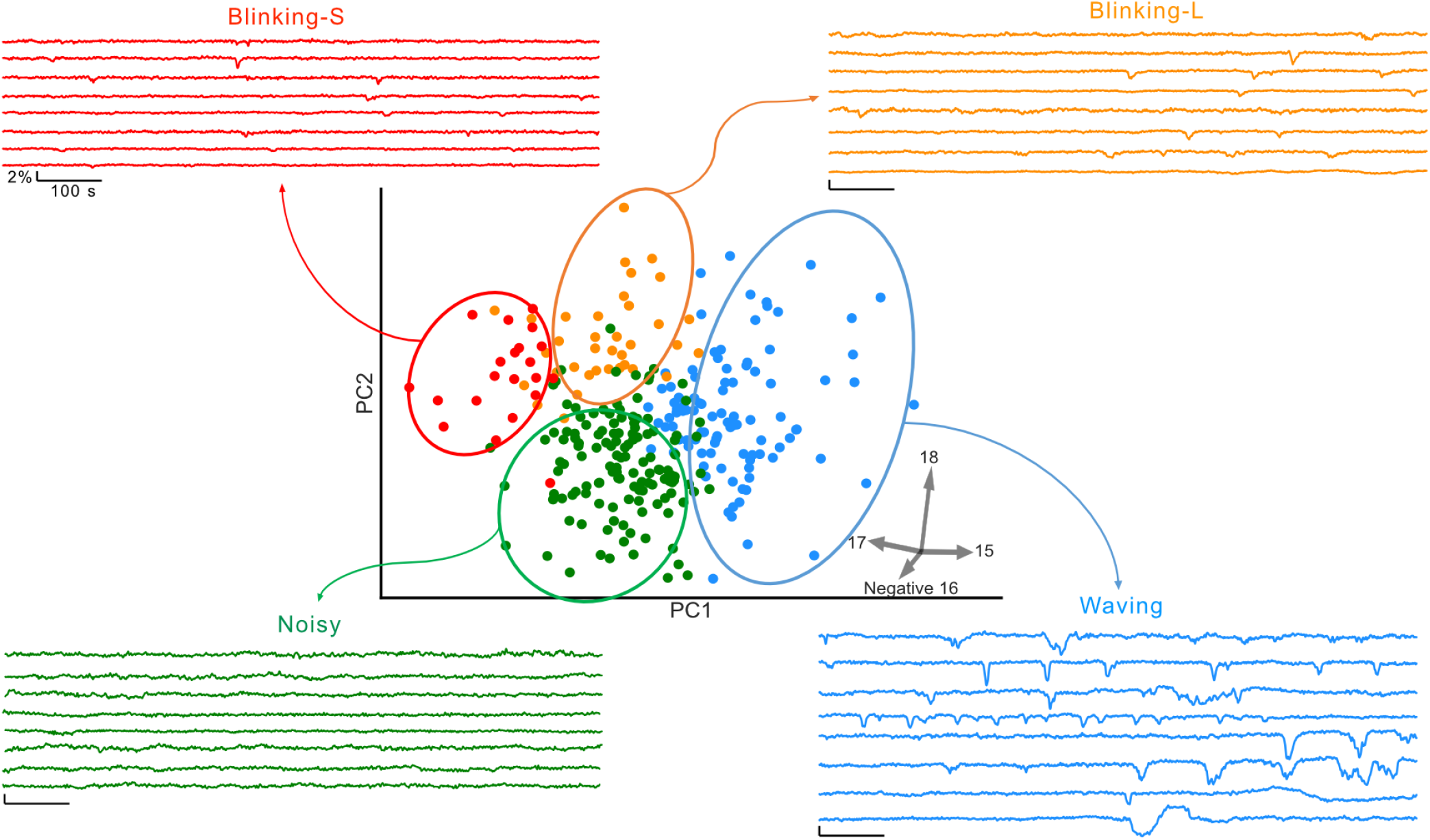
Exemplar time series for each DES class. Representative time series were selected by sorting the values of the feature’s pointing in the direction of each type in 2D PC space.

### TTX decreases *V*_*m*_ fluctuations in MDA-MB-231 cells

To assess the role of VGSC activity in the DES of MDA-MB-231 cells, we treated cultures with tetrodotoxin (TTX), a potent and specific inhibitor of VGSCs^14^. The ability of electrochromic dyes to report the effects of TTX is well established^32^. TTX decreased the frequency of *V*_*m*_ fluctuations, especially of large amplitude hyperpolarizations (-VEs, Figure 5A,B), in a dose-dependent manner. In particular, 10 µM TTX decreased the mean event rate by ~4x, from 9.5 × 10^−5^ to 1.97 × 10^−5^ events/cell/s (p < 10^−6^, one-sided bootstrapped significance test on the mean negative event rate per cell; Figure 5C). 1 µM TTX decreased the mean event rate by a lesser factor of ~2x, from 1.04 × 10^−4^ to 4.99 × 10^−5^ events/cell/s (p = 0.019, one-sided bootstrapped significance test on the mean negative event rate per cell; Figure 5D). The effect of TTX on event frequency recovered following washout (10 µM TTX; Figure 5E). For the feature-based analysis, projection of active TTX-treated MDA-MB-231 cells onto the active untreated cells’ PC space did not reveal DES class differentiation with TTX (Supplemental Figure S3).

**Figure 5.**
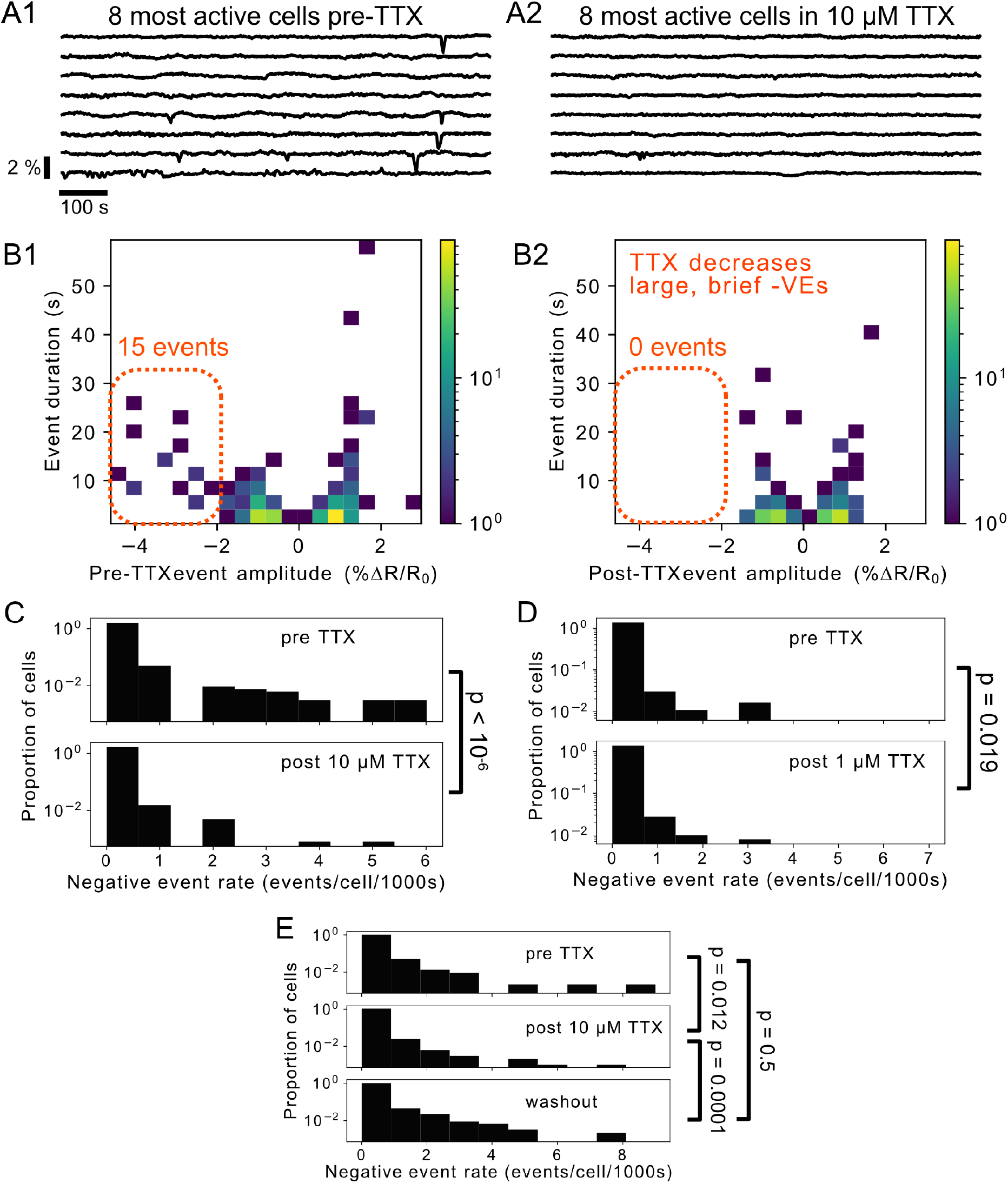
TTX decreased *V*_*m*_ fluctuations in MDA-MB-231 cells. (A1,2) Example *V*_*m*_ time courses from the 8 most active cells in acquisitions before (A1) and with (A2) 10 µM TTX. (B1, B2) Log-scaled 2D histograms of detected events before (B1) and with (B2) 10 µM TTX. TTX eradicates virtually all large, hyperpolarizing events (-VEs, dashed orange outline). (C) Log scaled histograms of the −VE event rate per cell for pre- and post-application of 10 µM TTX. The reduction in the mean from 9.5 ×10^−5^ events/cell/s to 1.97 ×10^−5^ events/cell/s (~4x decrease) is significant with p < 10^−6^. (D) The effect of TTX is dose dependent: 1 µM TTX reduced the mean event rate by a lesser factor of ~2x from 1.04 ×10^−4^ events/cell/s to 4.99 ×10^−5^ events/cell/s (p = 0.019). (E) The effect of TTX is reversible. The effects of TTX can be reversed by washing out the toxin, significantly increasing the −VE rate.

### *V*_*m*_ dynamics in MCF-10A cells

To assess the impact of MDA-MB-231’s cancerous, aggressive phenotype on *V*_*m*_ dynamics, we compared its *V*_*m*_ dynamics to that of non-tumorigenic MCF-10A cells (Figure 6A,C). Applying the same imaging protocol, we observed that only a small subset (0.46% ± 0.14%, n = 12 coverslips) of MCF-10A cells exhibited *V*_*m*_ fluctuations compared to MDA-MB-231 cells (6.84% ± 0.97%, n=22 coverslips, Figure 6E). Interestingly, incubation of the cells in TGF-β1 (5 ng/mL), a growth factor known to stimulate EMT, increased the percentage of cells exhibiting *V*_*m*_ fluctuations to 0.81% ± 0.19% (n = 13 coverslips, Figure 6B,D). With TGF-β, the mean −VE rate increased from 2.85 × 10^−6^ to 1.4 × 10^−5^ events/cell/s (p = 0.000567, one sided bootstrap difference of means at the cell level, Figure 6C-E).

**Figure 6.**
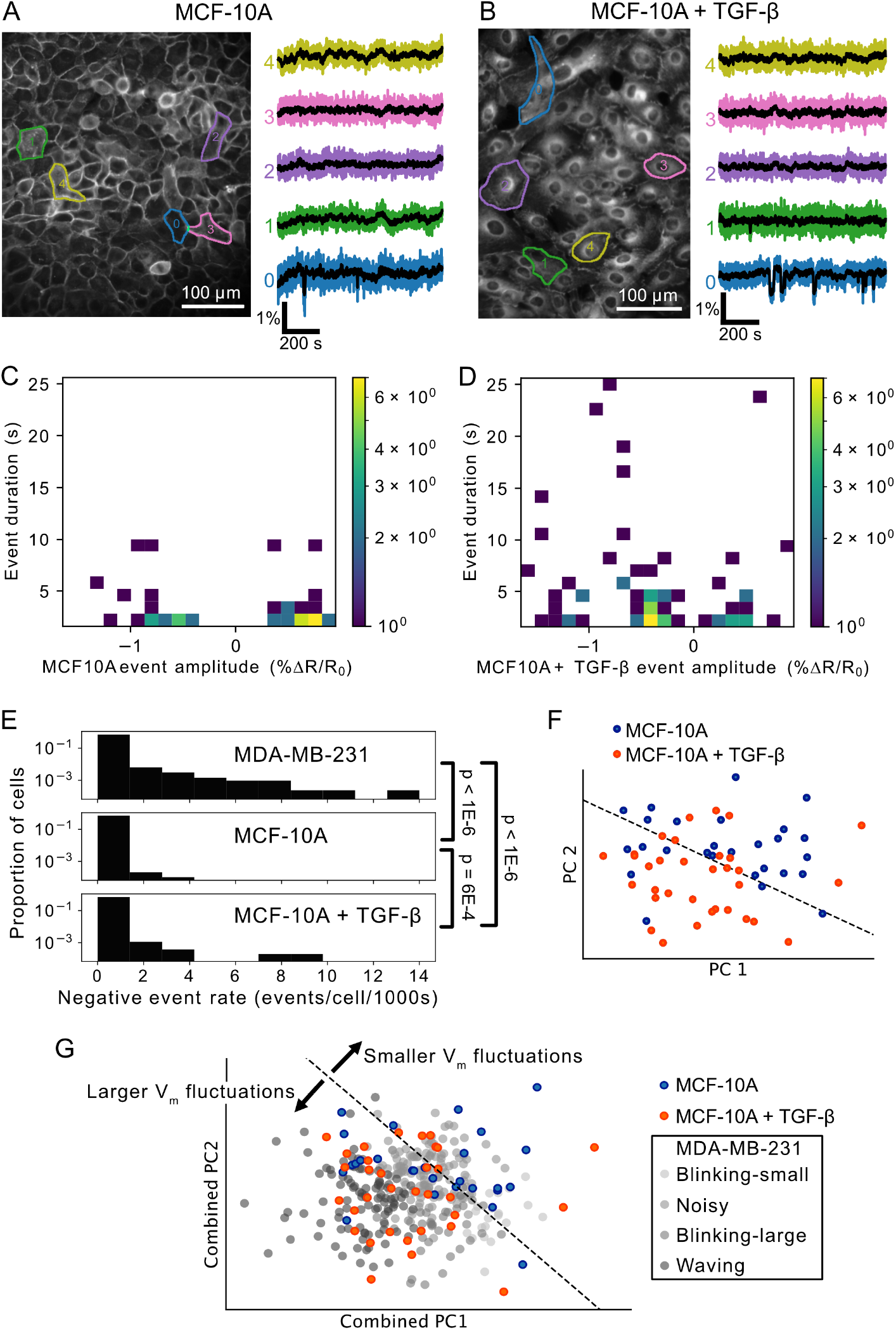
Non-cancerous MCF10A cells exhibit minimal *V*_*m*_ fluctuations which are upregulated by TGF-β. (A & B) Images and cellular *V*_*m*_ time series extracted from MCF-10A cells (A) and MCF-10A treated with TGF-β (B). (C & D) 2D log-scaled amplitude-duration histograms of (C) MCF10A and (D) MCF10A+TGF-β events. Both MCF-10A and MCF10A+TGF-β exhibit significantly lower event rates than MDA-MB-231 cells. E) Cell-level comparison of the negative event rate. Incubation of cells with TGF-β significantly increases the event rate. (F) Incubation of MCF-10A in TGF-β alters the cellular-level DES. The treated and untreated cells locate to different areas of 2D PC space, indicating DES differentiation between these groups. The black dashed line is the best linear separator calculated on the TGF-β treated (orange) and untreated (blue) groups. (G) Unlike the untreated MCF-10A which exhibited *V*_*m*_ fluctuations of varying amplitudes, TGF-β treatment biased active MCF-10A *V*_*m*_ fluctuations such that they localized near the large-amplitude “waving” MDA-MB-231 class in combined PC space.

Visualization of the MCF-10A cells’ normalized features in 2D PC space shows separation of DESs between TGF-β-treated and untreated MCF-10A cells (Figure 6F). In particular, 7/27 untreated MCF-10A cells and 9/33 TGF-β treated MCF-10A cells fell outside of the decision boundary of the best linear separator (Figure 6F, dashed line). In a feature space combining MDA-MB-231 DESs and MCF-10A DESs, the MCF-10A DESs co-localized over a range of MDA-MB-231 DES classes in 2D PC space, while TGF-β treatment increased localization with large-amplitude *V*_*m*_ “waving” MDA-MB-231 cells in the combined PC space (Figure 6G). In summary, TGF-β treatment increased the similarity of MCF-10A *V*_*m*_ dynamics with the large-amplitude “waving” MDA-MB-231 DES class.

### MDA-MB-231 cells *V*_*m*_ event synchrony

We observed temporal correlations between events occurring in simultaneously imaged MDA-MB-231 cells. Pairwise comparison of cellular *V*_*m*_ time series revealed a mixture of synchronous and asynchronous events (Figure 7A), indicating that these temporal correlations were not caused by optical crosstalk. To assess whether the *V*_*m*_ event temporal correlations occurred at rates significantly above chance, we generated event rasters for *V*_*m*_ transients detected in time bins ranging from 1-100 seconds (Figure 7B, showing 10 second bins). We then calculated the pairwise Pearson correlation coefficient (PCC) for all cells in each FOV. We compared the mean PCC for the cells to the PCC of randomly shuffled rasters, which preserved the cellular event statistics but destroyed any inter-cell temporal correlations. PCC increased overall with bin size as expected, and the PCC for the real event rasters was significantly greater than the PCC of the scrambled rasters for all bins sizes (Figure 7C). This result indicates a significant inter-cell temporal correlation of MDA-MB-231 *V*_*m*_ events, above that which would be observed by chance for such events randomly distributed in time.

**Figure 7.**
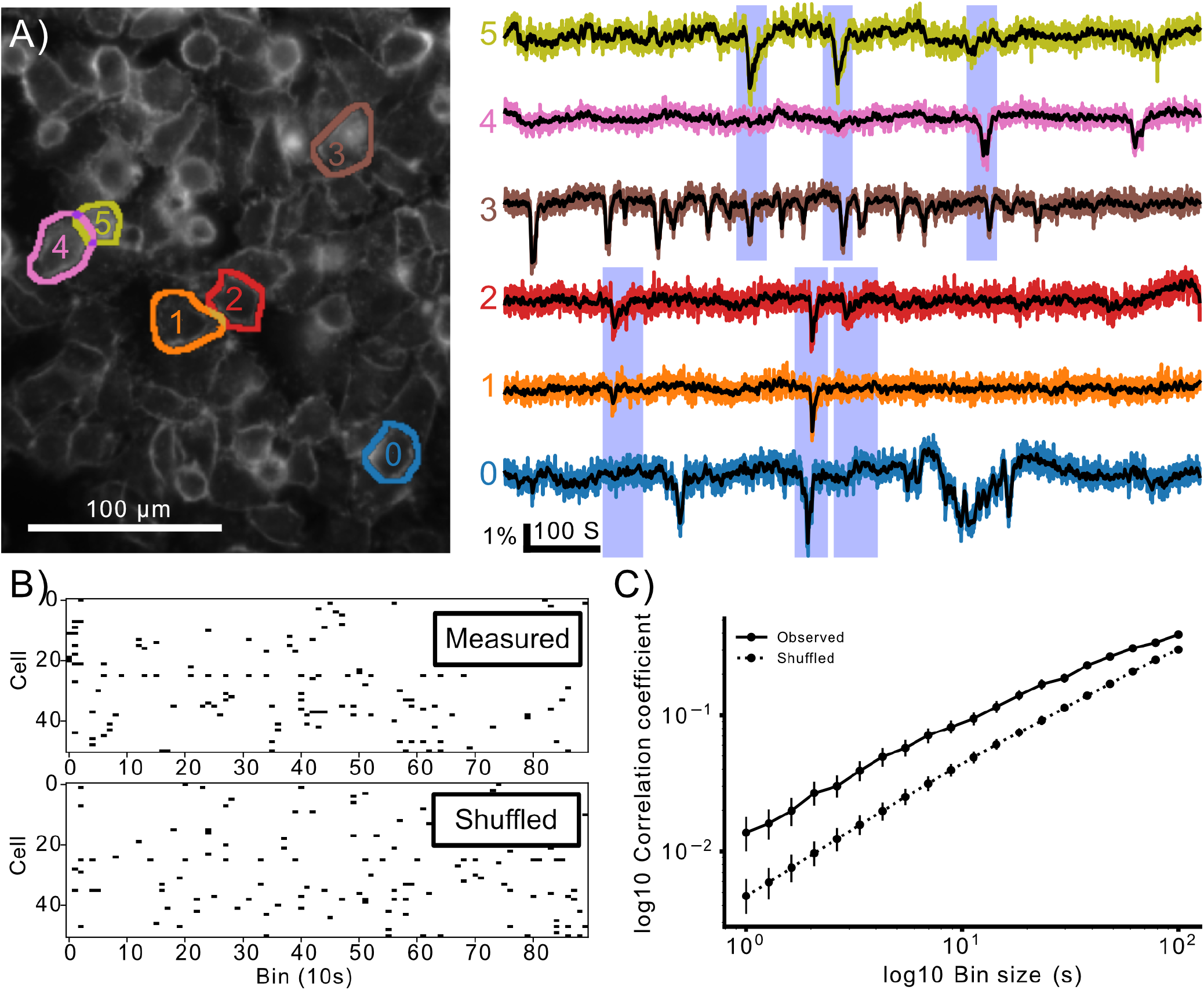
Transient activity is temporally correlated in MDA-MB-231 cells. (A) Example of temporally correlated cellular time courses from one FOV. Spatially separated cells (0,1,2) and (3,4,5) display synchronised hyperpolarisations. Importantly these synchronisations occur in different subsets of the groups and in spatially separated cells, indicating the synchronisation is not simply due to cross-talk in the imaging. (B) Quantification of correlation. Events are put into time bins, generating cellular event rasters. The average pairwise correlation between cells in a recording was calculated. A null hypothesis of zero correlation with the same temporal statistics was generated by temporally shuffling each cell’s event raster to obtain an estimate of the expected level of correlation if there were no cellular synchronisation. (C) The observed pairwise correlations are significantly higher in the observed data than shuffled data for all bin sizes, indicating the cellular activity is temporally correlated. The points plot the mean, and bars indicate the 95% confidence interval for the measured and shuffled case for each bin size.

In one instance we observed a wave of transient depolarizations propagating through a subset of cells unidirectionally across the FOV (Figure 8). The slope of the line fit to the distance as a function of hyperpolarization peak time from the first active cell shows a propagation speed of 27 µm/s (Figure 8B, supplementary Movie M2).

**Figure 8.**
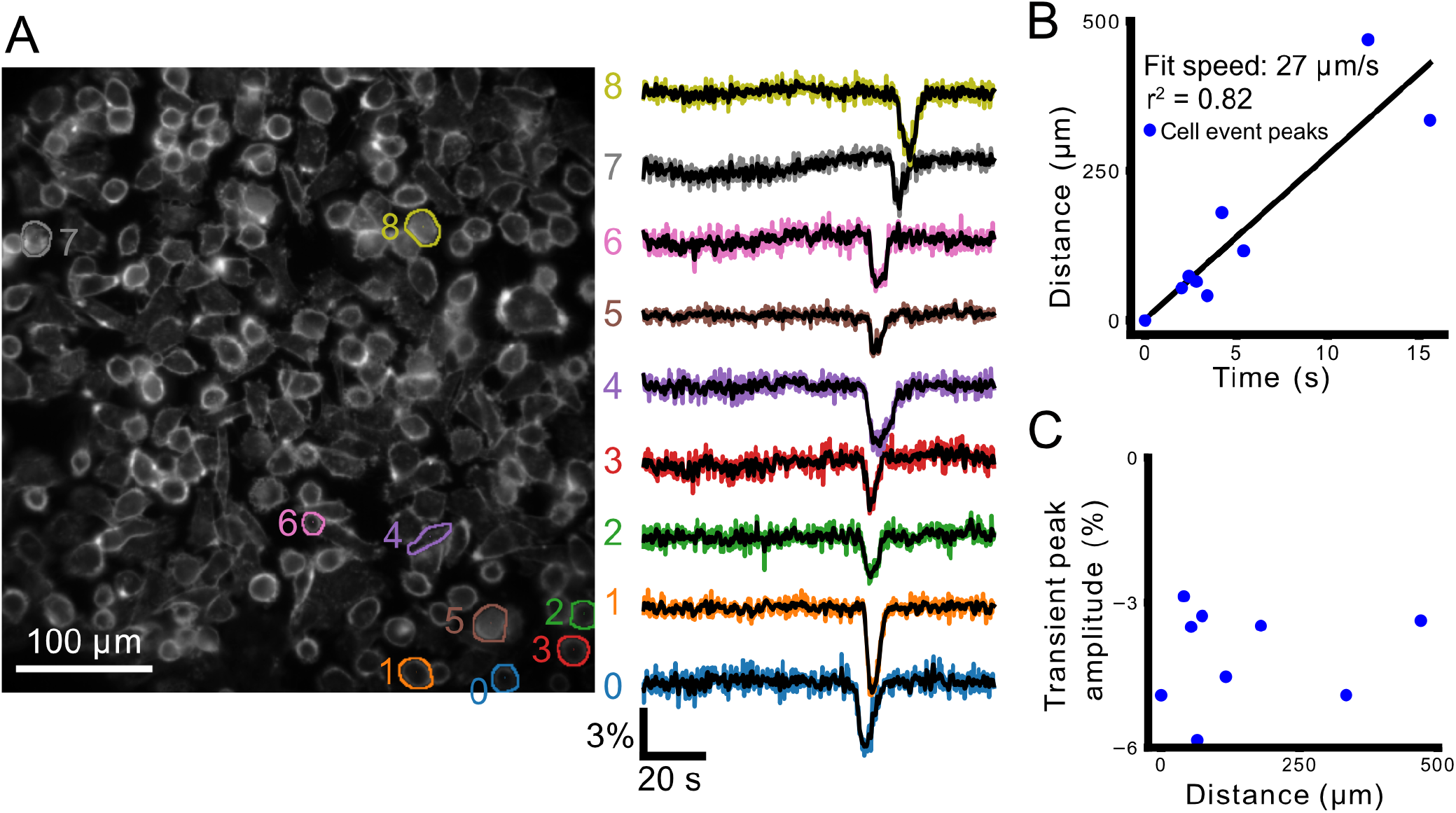
A wave of transient hyperpolarizations propagates across the FOV. (A) shows cellular ROIs and their color-corresponding Δ*R/R*_0_ time series (right). (B) plots distance from the first cell as a function of time at maximum hyperpolarization. The line fit to this relationship shows a propagation speed of 27 µm/s. (C) plots the peak transient amplitude (%Δ*R/R*_0_) as a function of distance from the cell first showing the transient hyperpolarization.

## Discussion

The high-throughout, cellular-resolution imaging data indicate that *V*_*m*_ fluctuates dynamically ~7% of highly aggressive, triple-negative MDA-MB-231 breast cancer cells. Both the event-based and feature-based analyses indicated that the most stereotypical of these events were transient hyperpolarizing “blinks” and “waves.” These events featured significant inter-cellular temporal correlation. Application of TTX decreased the dynamic *V*_*m*_ activity in MDA-MB-231 cells. The *V*_*m*_ of non-cancerous MCF-10A breast epithelial cells was comparatively static. Treatment with TGF-β increased *V*_*m*_ fluctuations in the MCF-10A cells and increased the feature-based similarity of their temporal fluctuations to those of MDA-MB-231 cells.

Whole-cell patch-clamp is the gold-standard for absolute *V*_*m*_ measurement. This technique measured that the steady-state resting *V*_*m*_ of human breast cancer cells is strongly depolarized relative to normal epithelia^15^,^18–20.^ Relatively little, however, had been reported on spontaneous *V*_*m*_ dynamics in human breast cells. Whole-cell current-clamp could in principle detect *V*_*m*_ fluctuations, but this has not been reported to date. It is possible that dialysis of the intracellular space by the patch pipette washes out or dampens the molecular machinery underlying the *V*_*m*_ fluctuations. Moreover, the low throughput of patch-clamp recording could hinder detection of the heterogeneous DESs exhibited by only approximately 1 in 20 cells.

Our detection of *V*_*m*_ fluctuations in subsets of MDA-MB-231 and MCF-10A cells was enabled by the novel application and adaptation of electrochromic voltage dye imaging. Critically, our approach enabled monitoring of spontaneous *V*_*m*_ fluctuations in hundreds of cells simultaneously at single-cell resolution. Electrochromic dyes track *V*_*m*_ with sub-microsecond temporal fidelity^33^, but are also phototoxic, which limits the exposure duration, rate, and total imaging time (3 ms exposures at 5 Hz over 920 seconds in our case). In the future, utilization of probes based on other mechanisms such as photo-induced electron transfer^34^, or transfection with fluorescent protein-based genetically-encoded voltage indicators (GEVIs)^35^, may increase the total imaging time and/or rate. In the case of GEVIs, however, photon budget is often limited by photobleaching, a problem addressed by newly-developed chemigenetic sensors^36^. Our ratiometric excitation scheme reduced imaging artefacts due to photobleaching, and variations in concentration and volume. It is important to note that voltage dye intensity reports relative changes in *V*_*m*_, not absolute *V*_*m*_. A second limitation of our analysis is that to remove the effects of bleaching and cell movement, we temporally high pass filter our *V*_*m*_ traces before analysis via division by the 1000 point rolling average of the time course. This filter precluded detection of activity varying on timescales slower than approximately 0.01 Hz.

A spiking, TTX-sensitive *V*_*m*_ phenotype was recently described in MDA-MB-231 cells through extracellular multi-electrode array (MEA) recording^28^. In this study, each large area (2 mm^2^) electrode aggregated signals from 100s of cells, enabling the detection of rare spiking events from pooled populations. While our 5 Hz imaging rate could not capture the “fast spiking” (lasting 10s of milliseconds) activity described by this study, it is possible that the “square-shaped” pulses (lasting up to 7 seconds) the authors detected arise from the same *V*_*m*_ dynamics reported as blinking/waving in our *V*_*m*_ image series. Our imaging methodology complements high temporal-resolution MEA recordings by enabling, for the first time, high throughput cellular-resolution localization of *V*_*m*_ fluctuations, necessary to detect spatiotemporal patterns, e.g., intercellular *V*_*m*_ wave propagation (Figure 8). The high throughput of *V*_*m*_ imaging demonstrated here opens a new window onto the ‘electro-excitable’ pathophysiology of cancer.

Roughly 7% of MDA-MB-231 cells demonstrated transient fluctuations in *V*_*m*_. The large amplitude events were hyperpolarizing ‘blinks’ and ‘waves’. This observation appears in stark contrast to classically excitable tissues like brain, heart, and muscle that have a resting *V*_*m*_ close to the reversal potential for K^+^, and which depolarize through the conductance Na^+^ and/or Ca^2+^ when excited^14^. Resting MDA-MB-231 cells exhibit a relatively depolarized *V*_*m*_, owing at least in part to a Na^+^ ‘window current’ conducted by Nav1.5^19,25,37^. The large negative-going events imply that K^+^ (possibly Cl^−^) conductance transiently increases, or that Na^+^ conductance transiently decreases, hyperpolarizing *V*_*m*_ toward the Nernst potential of K^+^. Moreover, the transient hyperpolarizations could be coupled to Ca^2+^ fluctuations^38^ through Ca^2+^-activated K^+^ channels, e.g. KCa3.1 preferentially expressed in metastatic breast tumors^39^. Future studies can elucidate the interplay between Ca^2+^ and *V*_*m*_ through simultaneous calcium and voltage imaging, and by pharmacological or genetic manipulation of KCa conductance during *V*_*m*_ imaging.

TTX drastically reduced hyperpolarizing transient (-VE) frequency in MDA-MB-231 cells. In the presence of TTX, blockade of persistent Na^+^ currents hyperpolarizes resting *V*_*m*_ in MDA-MB-231 cells^37^. Resting *V*_*m*_ hyperpolarization would decrease the driving force for the conductance of the transient hyperpolarizing current which could account, at least partially, for the reduction of transient hyperpolarizations we detected in the presence of TTX. Further work with e.g. ion channel pharmacological agents and gene silencing is required to elucidate the channels and currents mediating the transient hyperpolarizations observed here. Such experiments require the measurement of absolute membrane potential, which could be enabled optically through *V*_*m*_ fluorescence lifetime imaging (VF-FLIM)^40,41^. However, current VF-FLIM implementations have a temporal resolution <0.5 Hz and therefore unable to track the rapid fluctuations characterized here.

The heterogeneity of the *V*_*m*_ fluctuations concurs with the heterogeneous VGSC expression in these cells^15^. The largest amplitude fluctuations were transient hyperpolarizing “blinks” and “waves”. These hyperpolarizations could directly modulate the amplitude of the “window” current conducted by Nav1.5^19,25^, changing intracellular Na^+^ concentration and downstream Na^+^-modulated signaling pathways, e.g. SIK^17^. A primary function of the VGSC activity in these cells is controlling proteolysis via pericellular acidification driven by sodium-hydrogen exchange (NHE1)^42,43^. The fact that the hyperpolarizing *V*_*m*_-driven VGSC activity would occur intermittently in the cells would be suggestive of the following. First, it could ensure (i) better control of the proteolysis, i.e. invasion. Second, it would prevent excessive influx of Na^+^ into cells and possible osmotic imbalance that could compromise cell viability^44^. Further work is required to evaluate the potential mechanisms and consequences of *V*_*m*_ fluctuations.

Our results support the notion that fluctuating *V*_*m*_ is related to cancer cell aggressiveness. First, it was much more common in the strongly metastatic MDA-MB-231 cells compared with ‘normal’ breast epithelial MCF-10A cells. Second, blocking VGSC activity dampened the *V*_*m*_ fluctuations. Third, conversely, the dynamic activity of *V*_*m*_ in MCF-10A cells increased after treatment with TGF-β known to induce aggressive behaviour in these cells^45^. In the future, the *V*_*m*_ imaging can be exploited further to determine, at a cell-by-cell level, the correlative and causal relationships between *V*_*m*_ behavior and metastatic and structural phenotypes.

## Methods

### Cell culture

We cultured MDA-MB-231 cells in high glucose DMEM (Life technologies 41966029) supplemented with 5% FBS (Sigma, #F7524) and Penicillin-streptomycin (Sigma, #P4333). For imaging, we plated 10k – 30k cells on 12 mm collagen-coated glass coverslips (rat tail collagen, Sigma, #122-20). Cells were plated the afternoon prior to imaging. Imaging was performed in phenol red-free Liebovitz’s L-15 medium (Thermo fisher, #21083027), except during the high-K^+^ control experiments where mammalian physiological saline (MPS) was used instead (described below).

MCF-10A cells were obtained from ATCC. They were cultured in DMEM/F12 (GIBCO, #31331) supplemented with 5% Horse Serum (GIBCO, #16050), 10 µg/ml insulin (Sigma, #I-1882), 20 ng/ml Epidermal Growth Factor (Sigma, #E-9644), 100 ng/ml cholera toxin (Sigma, #C-8052), 500 ng/ml hydrocortisone (Sigma, #H-0888) and 100 mg/ml penicillin/ streptomycin (GIBCO, #15070). Cells were confirmed to be mycoplasma-negative (e-Myco Mycoplasma PCR Detection Kit, iNtRON Biotechnology). Passage was carried out using 0.25% trypsin-EDTA (GIBCO) followed by centrifugation (1000 rpm, 4 min) and resuspension in complete medium. Some sets of MCF-10A cells were cultured in Transforming Growth Factor-β1 (TGF-β; Peprotech, #100-21) at 5 ng/mL in complete media for 48 - 169 hours to drive EMT.

### Imaging

We prepared a 200 µM stock solution of the electrochromic voltage dye di-4-AN(F)EP(F)PTEA^29^ (100 nmol aliquots, Potentiometric Probes) in L-15 solution. Stock solution was kept for a maximum of 2 days after dissolving. Immediately prior to imaging, we gently washed the coverslip-adhered cells three times with warmed L-15 before placing them under the microscope. The coverslip was weighted in place with a tantalum ring and submerged in the dye diluted in L-15 to a final concentration of 3 µM. The cells rested in this configuration for 15 minutes before imaging. During imaging, cells were maintained at a temperature between 30C-37C by a home-made open-loop water perfusion system or by a closed-loop heated chamber platform (TC-324C, PM-1, Warner Instruments).

Our custom-built widefield epifluorescence microscope formed an image of the cells through a 25x 1.0 NA upright water dipping objective (XLPLN25XSVMP, Olympus) and 180 mm focal length tube lens (TTL180-A) onto a scientific complementary metal-oxide-semiconductor (sCMOS) camera (Orca Flash 4 v2, Hamamatsu). Imaging was performed with two-color sequential excitation and imaged in a single spectral channel. Fluorescence was ratiometrically excited in two channels resulting in opposite direction voltage signals in the collected emission (Figure 1C). LEDs were driven using a Cairn OptoLed (P1110/002/000). The first channel illuminated with 405 nm LED (Cairn P1105/405/LED), filtered with a 405/10 nm bandpass (Semrock LD01-405/10), and combined with a 495 nm long pass dichroic (Semrock FF495-DI03) and an additional 496 nm long pass (Semrock FF01-496/LP). The second channel illuminated with a 530 nm LED (Cairn P1105/530/LED), filtered with 520/35 nm filter (Semrock FF01-520/35), and combined with a 562 nm longpass dichroic (Semrock FF562-DI03). Emission was collected through a 650/150 nm bandpass filter (Semrock FF01-650/150) onto the sCMOS camera (Figure 1A). Images were acquired in Micromanager 2^46^ with the Orca Flash 4’s ‘slow scan’ mode, using the global shutter and frame reset with 4×4 digital binning. Imaging was performed at 5 Hz. During every image period, a 3-millisecond-exposure frame illuminated with each LED was acquired in rapid succession (Supplemental Figure S1). Illumination intensities for each channel were approximately matched between each channel and adjusted to give a signal intensity of around 4000 counts/pixel in labelled cell membranes. Intensities were typically between 0.1 and 1.3 mW/mm^2^ for the blue excitation and 1.5 and 3.4 mW/mm^2^ for the green excitation.

### Imaging Protocols

We acquired each trial, consisting of sequences of 10,000 frames (5,000 dual color-excited acquisitions) at different locations on the coverslip. Imaging locations were selected from confluent areas (median 975 cells/mm^2^, interquartile range 605, 1247 cells / mm^2^). We acquired between 1 and 6 trials per coverslip, with each trial occurring in a distinct location. In tetrodotoxin (TTX) experiments, we first imaged 1-2 control trials without TTX (‘pre’ trials). We then added 1 mM of TTX citrate (Abcam, ab120055) in PBS stock solution to the imaging medium to achieve the a final concentration of 1 or 10 µM. 1-4 trials were imaged in the presence of TTX (‘post’ trials). For certain 10 µM TTX experiments, following 1-2 trials acquired in the presence of TTX, we replaced the TTX-containing medium with regular dye-containing L-15 medium and imaged 1-2 trials in this condition (‘washout’).

High potassium wash-in trials were conducted in mammalian physiological saline (MPS^38^), consisting of (in mM): 144 NaCl, 5.4 KCl, 5.6 D-Glucose, 5 HEPES, 1 MgCl_2_, 2.5 CaCl_2_. Mid-trial, a high-potassium solution was washed in to depolarise the cells for validation of the voltage dye function. This solution was osmotically balanced consisting of (in mM): 49.4 NaCl, 100 KCl, 5.6 D-Glucose, 5 HEPES, 1 MgCl_2_, 2.5 CaCl_2_.

### Patch-clamp voltage dye calibration

We assessed the range of fluorescence change expected for known changes in *V*_*m*_ through whole-cell voltage-clamp and simultaneous voltage imaging. Cells were imaged in phenol red-free Liebovitz’s L-15 medium at room temperature. Healthy, dye-labelled cells were selected and patched with pipettes between 3 and 10 MΩ. The pipette contained an Ag/AgCl bathed in intracellular solution (in mM): 130 K-Gluconate, 7 KCl, 4 ATP - Mg, 0.3 GTP - Na, 10 Phosphocreatine - Na, 10 HEPES. Voltage-clamp signals were amplified with a Multiclamp 700B (Molecular Devices) and digitized with Power 1401 (Cambridge Electronic Design) using Spike2 version X.

Ratiometric imaging was performed as described above, but at an increased rate of 100 frames/second. During each imaging trial, *V*_*m*_ was clamped for 1 s epochs at values varying between –60 to +30 mV in 10 mV increments. Fluorescent time courses were extracted from a cellular region of interest (ROI) around the patched cell for both excitation channels. The trials were bleach-corrected and converted to Δ*F/F*_0_ using a linear fit to their time course. We calculated the average blue to green excited frame ratio (Δ*R/R*_0_) across trials at each holding potential. A line was fit to Δ*R/R*_0_ vs *V*_*m*_. The line gradient reflects the sensitivity of Δ*R/R*_0_ to *V*_*m*_ (% change per 100 mV) for each cell (n=12 cells).

### Image Processing

All data analysis was performed in Python 3 using NumPy^47^, SciPy^48^, Tifffile, Scikit-image^49^, Scikit-learn^50^ and Pandas^51^. Figures were generated using MatPlotLib^52^. The analysis code is available at https://github.com/peq10/cancer_vsd. Our dual-color excitation scheme generated image time series interleaving blue and green light-excited frames (Figure 2A). We subtracted the constant dark value from each frame and separated the time series into two color channels. We applied a pixel-wise high pass filter, rejecting signals slower than 0.01 Hz, to the separated time series. In particular, slowly varying signals (mainly bleaching) were removed from each channel by dividing the stacks pixel-wise by a temporally filtered version of themselves. The filter was a uniform filter of length 1000 points. The filter was symmetric (i.e. time point t0 affected by points t>t0 and t<t0 equally). These filter-normalised time series were then divided, blue frames by green frames, to find the ratio image for each time point (Figure 2A). Cells were segmented using CellPose^53^, using the default cytoplasmic segmentation model and approximate cellular diameter of 30 pixels. Additional segmentations of active cells that the Cellpose network did not identify were added by hand. The segmented ROIs were eroded with a single (1-round of binary) before extracting the ROI time courses to suppress the effects of movement at the cell edges. For each eroded ROI, we calculated the median of the pixel values at each time point. The mean of the time courses was subtracted and offset so that they were symmetric about 1.

### Event detection

We implemented an event detection algorithm to identify significant changes to Δ*R/R*_0_ reflecting the fluctuation of *V*_*m*_. We first calculated the time course of the intra-ROI pixel-wise standard deviation for each eroded ROI. We filtered the median (calculated as above) and standard deviation time courses with a *σ* = 3 point (i.e. 0.6 s) Gaussian filter. *V*_*m*_ fluctuation events were identified when the temporally filtered median pixel value diverged from 1, its time average value, by more than 2.5 times the temporally filtered standard deviation (Figure 1H). Short events were removed and neighbouring events merged by 2 iterations of binary opening and then 2 rounds of binary closing on the detected event array. Where events consisted of positive- and negative-going Δ*R/R*_0_, they were split into entirely positive going and entirely negative going events.

### Time series and event inclusion criteria

Cells were considered dying/dead and excluded from analysis where the raw pixel values increased in brightness by more than 25% during the acquisition in the blue channel, indicating loss of membrane polarization. Events arising from non-voltage related changes in brightness were identified as simultaneous events and excluded from the analysis. In the MCF-10A image series, where events were very rare, imaging artefacts were identified and excluded where more than 3 events overlapped by more than 30% in time. In MDA-MB-231 data, events were excluded where more than 5 events overlapped by more than 50% in time. Following event detection, all active cell time series were then evaluated by visual inspection of processed videos to reject events caused by floating dust, focal shifts, or other apparent imaging artefacts. Events satisfying both the automated and manual quality-control measures were analysed for event frequency, polarity, amplitude and duration.

In the feature-based analysis, we only included trials that completed the full 920 s imaging period (18279 out of 18752), and then applied the event detection algorithm described above to identify “active” cells (982 out of 18279). We then excluded all time series containing apparent imaging artefacts (dust, etc.). Following quality control and event detection, 297 MDA-MB-231, 28 MCF-10A, and 33 MCF-10A+TGF-β time series were admitted to the feature-based analysis pipeline described below (Figure 3A,B).

### *V*_*m*_ time series clustering analysis

Complementing the event-based analysis, developed the Cellular DES Pipeline to classify the ROI-extracted time series according to its most salient dynamic features extracted by the Catch-22 algorithm^31^. The analysis realized with the Cellular DES Pipeline provides insight into the time series characteristics beyond simple event detection and quantification, enabling classification of the heterogeneous *V*_*m*_ dynamics into like clusters.

From the admitted time series, we extracted 22 features from each cellular ROI’s median time series with the Catch-22 algorithm (Figure 3C). After plotting the distribution of the individual features, around 80% (259/324) of cells shared the same value for the feature corresponding to the first minimum of the automutual information function, which we subsequently excluded from the feature list. We rescaled the raw feature values for the remaining 21 features between 0.0001 and 1 and applied the Box Cox transformation to normalize their distributions. We then rescaled the normalized values to between 0 and 1 to ensure equal weighting into the clustering algorithms. To evaluate the number of Dynamic Electrical Signature (DES) classes, we implemented hierarchical clustering and Gaussian Mixture Modeling (GMM) on the 21 normalized features. Both clustering algorithms generated clusters with silhouette coefficients, which measure subtype dissimilarity^54^, decreasing to their lowest levels between 5 and 6 clusters (Figure 3D). Based on these silhouette scores and on visual inspection of the time series, we chose to sort the time series into four types as this resulted in the most homogeneous classes. Based on the general pattern of each type, we named the DES classes: small blinking (blinking-s), waving, noisy and large blinking (blinking-l).

To select exemplar time-series from each DES class (Figure 4), we performed Principal components analysis (PCA) and visualized the four classes in a 2-dimensional feature space. Each feature cluster occupies a unique area of the PC space. To identify exemplar time series of each type, we calculated the components of each feature and drew a vector for each feature’s coefficients of PC1 and PC2 (Figure 4). These vectors therefore point to the type exhibiting the corresponding features most saliently. To identify exemplar time series from each type, we sorted the time series according to the feature whose vector points to that type.

## Data Availability

The imaging and electrophysiological datasets generated and analysed during the current study are available from the corresponding author on reasonable request.

## Acknowledgements

The authors would like to thank Dr. Julia Gallinaro for her advice on the quantification of synchrony within the time courses. This work was supported by a Pump-prime Award from the Integrated Biological Imaging Network (IBIN3AF), the Royal Academy of Engineering under the RAEng Research Fellowships scheme (RF1415/14/26), the Biotechnology and Biology Research Council (BB/R009007/1), a Wellcome Trust Seed Award (Grant No. 201964/Z/16/Z), the Engineering and Physical Sciences Research Council (Grant No. EP/L016737/1 to Imperial College), and by a Cancer Research UK and Stand Up to Cancer UK Programme Foundation Award to CB (C37275/1A20146).

## Author contributions statement

PQ devised the image acquisition system and methods with the guidance from CDA and AJF. PQ, MBAD, CB, and AJF designed the experiments. PQ performed the experiments, analysed the images, and designed and performed the event based analysis. MAG cultured the MCF-10A cells. YS developed the Cellular DES Pipline with the guidance of CB, MBAD, AJF, and PQ. YS performed the feature-based analysis with assistance from PQ. PQ, YS, MAG, CB, MBAD, and AJF wrote the manuscript. All authors reviewed the manuscript.

