## Supplementary Figures for "Membrane voltage fluctuations in human breast cancer cells"

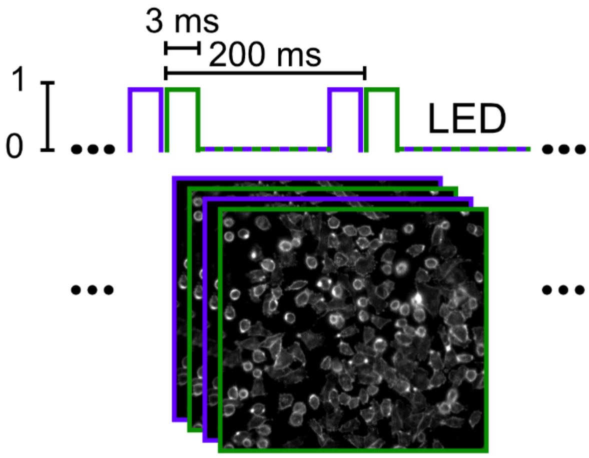


**Figure S1. Image acquisition timing diagram.** Top, illustration of the command signals sent to the LED illumination driver showing the length and duty cycle of the ratiometric illumination. Each color was sequentially strobed for 3 ms every 200 ms resulting in an interleaved series of fluorescent images (bottom).

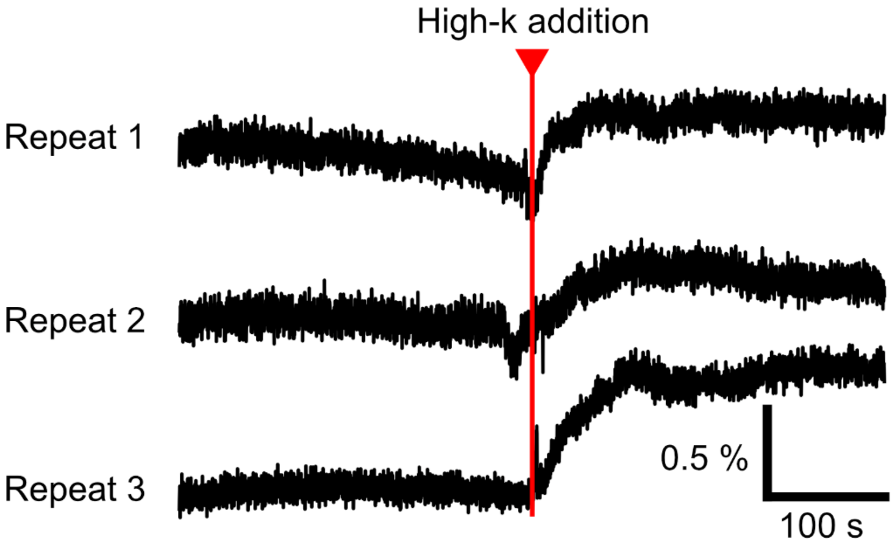


**Figure S2. Perfusion of cells with a high potassium solution results in global depolarisation.** Each trace shows the mean time course of all cells in the FOV during the perfusion of high potassium (containing 100 mM KCl) mammalian physiological saline. The resulting membrane depolarisation can be clearly observed in the voltage dye signal.

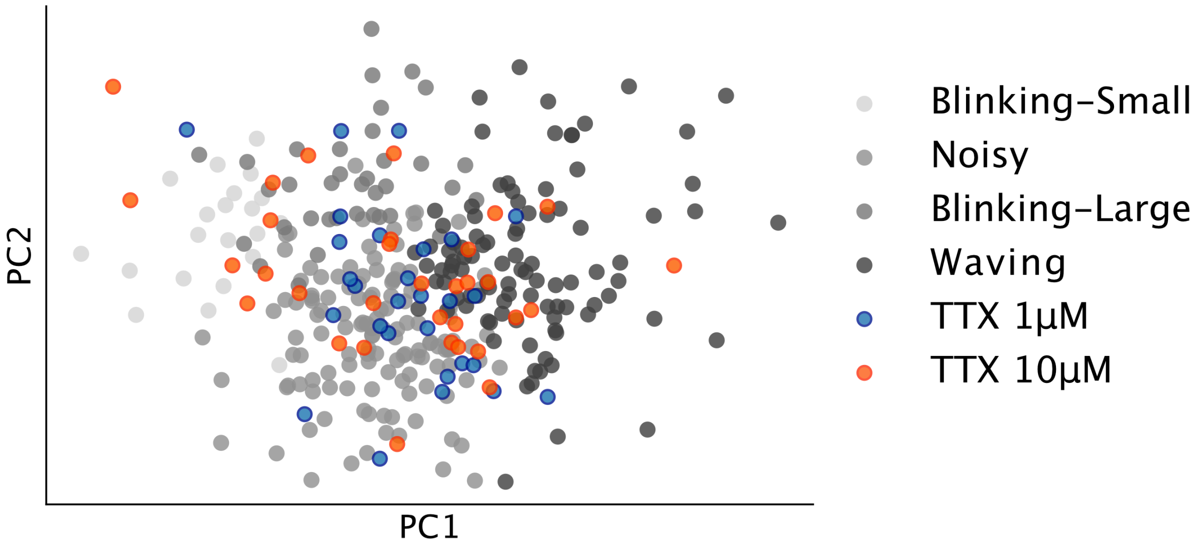


**Figure S3. Treatment of MDA-MB-231 cells with TTX did not bias their Vm fluctuations with respect to the four DES classes.** The projection of TTX-treated MDA-MB-231 cells (blue: 1 μM; orange: 10 μM) onto the untreated cells' 2D PC space shows distribution throughout the four DES classes (grey).

**Movie Legends:**

**Movie M1. Exemplar video of voltage transients in MDA-MB-231 cells.** Left, raw video of the blue fluorescent channel showing the voltage dye-stained cells. Right, video of the ratiometric pixel-wise $\Delta R/R_0$ calculated using the process described in the Methods. The video is shown at 25 frames/s (5x real speed). Darker colours correspond to hyperpolarisations and lighter colours to depolariastions. Large and short hyperpolarisations are the most prominent transients, and most cells display little or no voltage activity. This video has been spatially and temporally filtered with a gaussian of width (3,2,2) samples in (t,y,x) and the 0.5 and 99.5% percentile pixels are saturated to facilitate transient visualisation.

**Movie M2. Video of voltage transient wave.** Left, raw image of the blue channel fluorescence. Right, video of the ratiometric pixel-wise $\Delta R/R_0$ calculated using the process described in the methods. A wave of negative transients can be seen propagating from cells in the lower right corner to the upper left corner at an approximate speed of 27 μm/s. The video is shown at 25 frames/s (5x real speed). Darker colours correspond to hyperpolarisations and lighter colours to depolarisations. This video has been spatio-temporally filtered with a gaussian of width (3,2,2) samples in (t,y,x) and the 0.5 and 99.5% percentile pixels are saturated to facilitate transient visualisation.
